# Tracking malaria parasite lineages through *de novo* mutations in highly related *Plasmodium falciparum* genomes

**DOI:** 10.64898/2026.05.08.723589

**Authors:** Balotin Fogang, Marc-Antoine Guery, Ian Cheeseman, David J. Conway, Antoine Claessens

**Author notes:** equal contributors (Balotin Fogang, Marc-Antoine Guery).

## Abstract

Where malaria transmission declines, the remaining infections are increasingly low-density and asymptomatic, forming a persistent reservoir that is difficult to track using conventional epidemiological approaches. However, genomic data from such community-level infections remain scarce, limiting the ability to track parasite lineages, detect clonal expansions, and identify persistent chronic infections in pre-elimination settings. Here, 78 single-genotype *P. falciparum* genome sequences are analysed from community infections within a small area of The Gambia, where malaria transmission has substantially declined over recent decades. Pairwise identity-by-descent (IBD) analysis revealed generally low genetic relatedness among parasites, consistent with ongoing recombination and genetic mixing at the community scale. Nevertheless, eight clusters of near-identical genomes (IBD > 0.9) were identified, enabling the inference of recent *de novo* mutations that differentiate these genomes. Across these clusters, 43 *de novo* single-nucleotide polymorphisms and 19 short indels were identified using long-read-derived reference genomes. The observed pattern of mutation in natural infections broadly resembled that previously reported from laboratory mutation-accumulation experiments, including a strong transition bias and enrichment of G:C→A:T substitutions. These results demonstrate that combining IBD analysis with de novo mutation detection enables fine-scale resolution of parasite relatedness and recent transmission history. As malaria transmission continues to decline, such approaches may become increasingly valuable for tracking local transmission, identify parasite lineages, and potentially distinguish persistent infections from reintroduction events.

## INTRODUCTION

Malaria remains one of the most significant public health challenges worldwide, caused primarily by *Plasmodium falciparum*, which accounts for almost all malaria-related mortality(World Health Organization, 2025). Current understanding of *P. falciparum* genomics is predominantly derived from studying samples of clinical cases and culture-adapted parasites, with relatively few genomes sequenced from community-level or asymptomatic infections (Nyarko & Claessens, 2021). However, asymptomatic carriers play a critical role in sustaining malaria transmission and maintaining reservoirs of genetic diversity, particularly in regions transitioning toward malaria elimination (Abdi et al., 2023; Mwesigwa et al., 2024). More genomic data from such infections may help to understand parasite population genetics and evolution (Amambua-Ngwa et al., 2026; Fola et al., 2024; Guo et al., 2024).

As malaria transmission declines, evidence of occasional clonal lineage expansions and reduced parasite recombination becomes more apparent (Auburn & Barry, 2017). However, distinguishing different genetic processes in parasite populations remains challenging, particularly in endemic areas where transmission rates are still moderate(Cabrera-Sosa et al., 2024; Daniels et al., 2015; Pringle et al., 2019). Where endemicity remains sufficient to sustain genetic diversity but is declining, novel analytical frameworks may be useful to help prepare for elimination.

Genotyping of limited numbers of loci has provided valuable insights into population structure and diversity (Auburn & Barry, 2017; Tessema et al., 2019).However, whole-genome sequencing (WGS) allows investigation of finer-scale relationships among parasite genomes. One powerful approach involves the analysis of genomic identity-by-descent (IBD), which can measure the proportion of sampled pairs of parasite genomes inherited from a recent common ancestor. IBD analysis can identify highly related parasites arising from recent transmission events, clonal expansion, or inbreeding, and has been widely used to infer recent shared ancestry in population genetics (Browning & Browning, 2013). However, when genomes are nearly identical, population genetic markers may still fail to distinguish between individual parasite lineages.

Identification of inferred *de novo* mutational changes accumulated during parasite replication provide an additional source of genetic variation that can differentiate otherwise identical genomes (Redmond et al., 2018). In microbial pathogens, mutation-based approaches have proven valuable for reconstructing transmission chains and inferring evolutionary timescales (Didelot et al., 2014; Eyre et al., 2013; Metsky et al., 2017). Although mutation accumulation has been characterised in *P. falciparum* under laboratory conditions, the spectrum of naturally occurring *de novo* mutations within natural infections also deserves attention.

Here, we combine whole-genome sequencing, IBD analysis, and inferred *de novo* mutation detection to investigate *P. falciparum* population structure in a community cohort from eastern The Gambia. Using a dataset of 78 single-genotype parasite genomes sampled across four neighbouring villages, we identify highly related parasite clusters and characterise the mutations that distinguish them. These results indicate a proof-of-principle that mutation-based genomic approaches might help resolve transmission patterns and parasite persistence in settings approaching malaria elimination.

## MATERIAL AND METHODS

### *P. falciparum* genome sequences from community infections

Illumina short-read genome sequencing of *P. falciparum* infections in a rural area of The Gambia was obtained as described previously (Guery et al., 2024). Briefly, participants were recruited in four neighbouring villages (Madina Samako: K, Njayel: J, Sendebu: P, and Karandaba: N) located within a 5 km radius in Upper River Region (URR), eastern Gambia (Fogang et al., 2024). From December 2014 to December 2016, a total of nine community surveys were conducted in these villages to detect *P. falciparum* infections by qPCR. Additionally, participants presenting with malaria symptoms during one transmission season (September to December 2016) were also included. Finally, in December 2016, 42 asymptomatic *P. falciparum* carriers were enrolled in a longitudinal cohort with monthly follow-up sampling until May 2017 or until spontaneous clearance of infection (Collins et al., 2022). In total, out of 522 *P. falciparum* positive blood samples, 199 high-quality genomes were generated with Illumina sequencing under protocols managed by the MalariaGen consortium(Abdel Hamid et al., 2023). To avoid sampling bias, only one genome per continuous infection was retained for further analysis. In addition, two *P. falciparum* genomes from isolates cloned in culture and sequenced using PacBio HiFi technology were assembled *de novo* and used for specific comparative analyses (Nyarko et al., 2025).

### Fws and IBD calculations

As previously described (Guery et al., 2024), parasite diversity within each isolate genome was estimated by the *F*ws metric based on allelic frequencies from genomic data (Manske et al., 2012). Genetic relatedness between single-genotype genomes (*F*ws > 0.95) was estimated with hmmIBD (Schaffner et al., 2018), also already described (Guery et al., 2024). Briefly, to accurately evaluate the genetic similarity between parasite isolates, we calculated pairwise mean posterior probabilities of identity-by-descent (IBD) between genomes using hmmIBD, a hidden Markov model–based method that infers shared ancestry from meiotic recombination patterns, assuming a *P. falciparum* recombination rate of 13.5 kb/cM (Miles et al., 2016; Schaffner et al., 2018). IBD inference relies on shared homozygous sites between pairs of samples, which are used to estimate the proportion of the genome inherited from a recent common ancestor.

To identify putative chromosomal regions under selection, we quantified the extent of IBD sharing across the population of genomes. For all genome pairs with low overall relatedness (IBD < 0.5), we extracted genomic intervals inferred to be in IBD by hmmIBD. Overlapping IBD segments across different genome pairs were then merged to define a set of non-redundant genomic intervals. For each interval, we counted the number of genome pairs in which that interval was observed and divided this count by the total number of genome pairs analysed. This yielded a measure of IBD coverage, defined as the proportion of genome pairs sharing a given genomic interval.

### Identification of *de novo* single-nucleotide polymorphisms (SNPs)

Inferred recent *de novo* mutations were identified by characterising SNPs or short insertions/deletions (indels) that distinguished otherwise nearly identical monoclonal parasite genomes. We first identified clusters of genomes with high pairwise identity-by-descent (IBD > 0.9), and labelled them as ‘clusters’. Within each cluster, genomes were compared pairwise.

SNP variant data were identified for these samples within the MalariaGEN *Plasmodium falciparum* v7 global dataset (Abdel Hamid et al., 2023). Highly related genomes were then compared pairwise to identify candidate mutations. For each pair of highly related genomes, *de novo* SNPs were defined as genomic positions showing a difference in within-sample allele frequency greater than 0.6 between two otherwise almost-identical infection samples, restricted to genomic regions inferred to be in IBD. Genomic regions not in IBD were excluded from the analysis (Supplementary Figure 3). All putative *de novo* SNPs were visually inspected and manually validated using the Integrative Genomics Viewer (IGV) (Robinson et al. 2023).

### Identification of *de novo* insertions and deletions (indels)

Due to the very high A:T richness of the *P. falciparum* genome, it is normally difficult to obtain accurate indel detection in diverse isolates when mapping short-read sequencing data to a single reference genome such as that of the 3D7 strain. To overcome this limitation, we used long-read genome assemblies for two Gambian parasite isolates to identify *de novo* indels differentiating genomes within two of the clusters of related infections.

These long-read assemblies were generated following culture adaptation and cloning of one isolate from genotype cluster 9 and one from cluster 12 (Nyarko et al., 2025). Illumina short-read data from all isolates with IBD regions belonging to these clusters were mapped to their respective related long-read reference assemblies. Short indels were then identified using the Genome Analysis Toolkit GATK (McKenna et al., 2010) following the MalariaGen Pf7 pipeline process (MalariaGen et al. 2023). For each cluster, indel calls were filtered using stringent criteria to retain only high-confidence variants. Specifically, indels were excluded if they met any of the following conditions: Quality by Depth (QD) < 10.0, Depth of Coverage (DP) < 10, Fisher Strand Bias (FS) > 60.0, Symmetric Odds Ratio (SOR) > 3.0, Mapping Quality Rank Sum Test (MQRankSum) < −8.0, or Read Position Rank Sum Test (ReadPosRankSum) < −8.0.

Only indels located within the core genome, as defined by (Otto et al., 2018) and within chromosomal regions inferred to be in IBD were retained. In addition, indels located within homopolymer tracts longer than 20 bp were excluded, due to the known propensity of GATK to miscall variants in long homopolymer regions (Ross et al., 2013). All remaining indels were visually inspected and manually validated using IGV.

## RESULTS

### IBD analysis identifies genetically related *P. falciparum* genomes from community cases

Blood samples had been regularly collected between December 2014 and May 2017 from residents of four neighboring villages in eastern Gambia and tested for malaria parasites (Fogang et al., 2024). Out of 522 *P. falciparum* community infections detected, 199 infections were successfully genome sequencedusing Illumina short-read sequencing (Guery et al., 2024), of which 78 corresponded to high-quality unmixed single-genotype infections (within-host fixation index *F*_ws_ > 0.95) were retained for subsequent analyses (Figure 1, Table S1). To assess patterns of genomic relatedness among the different single-genotype *P. falciparum* infections, pairwise identity-by-descent (IBD) analysis was performed (Supplementary Figure 1). Among the 3003 pairwise comparisons (Table S2), 41 (0.21%) exhibited IBD values greater than 0.2, 32 (0.16%) exceeded an IBD threshold of 0.5, and 20 (0.67%) exceeded 0.9, indicating a generally low level of relatedness among circulating parasite genomes (Figure 2). Drug resistance markers can potentially skew the IBD due to selection reducing variation around each resistance locus, but this effect is negligible in the dataset (Supplementary Figure 2), and would not affect the principle of the analysis being performed here.

**Fig 1.**
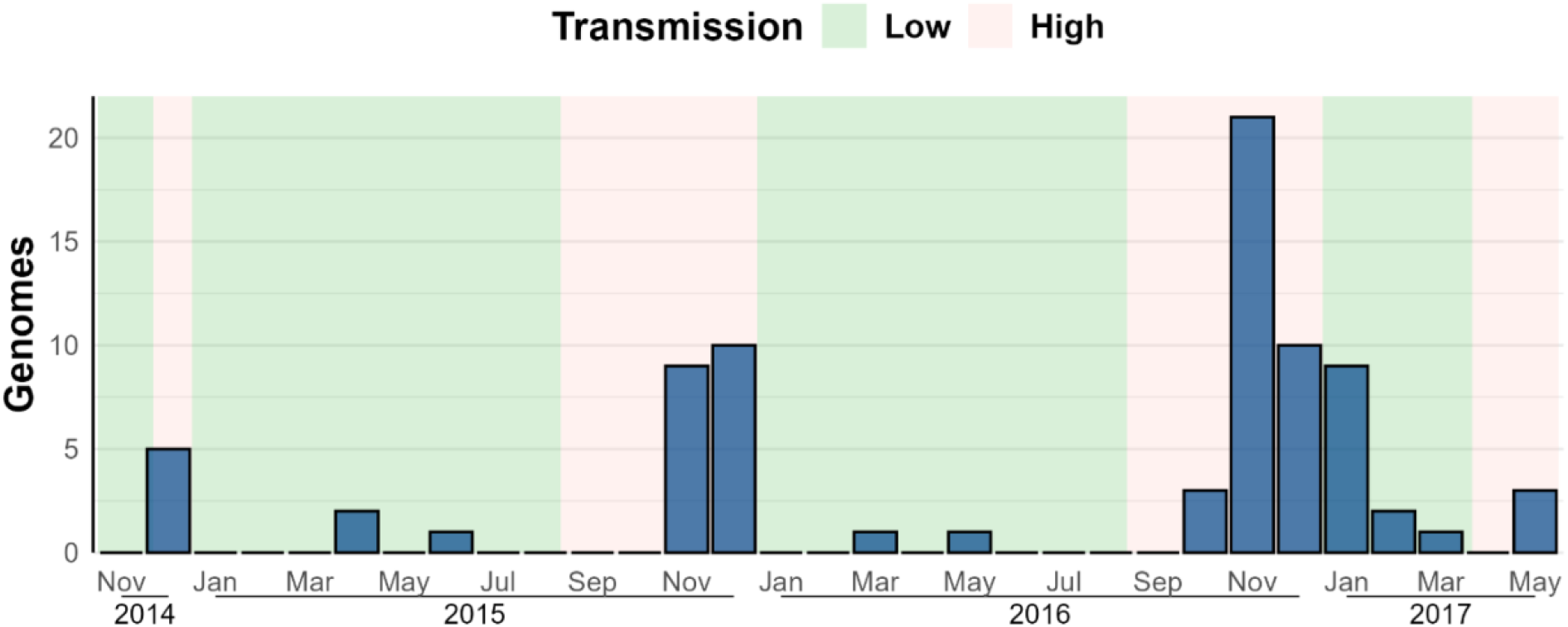
Timing of sampling of 78 *P. falciparum* monoclonal genomes from community infections. A total of 522 *P. falciparum*-positive blood samples were collected from participants living in four neighbouring villages in The Gambia. From these samples, 199 high-quality genomes were generated. To avoid repeated sampling of the same infection, only one genome per continuous infection was retained for further analysis. Shown here are the 78 genomes with Fws >0.95, classified as monoclonal infections.

**Fig 2.**
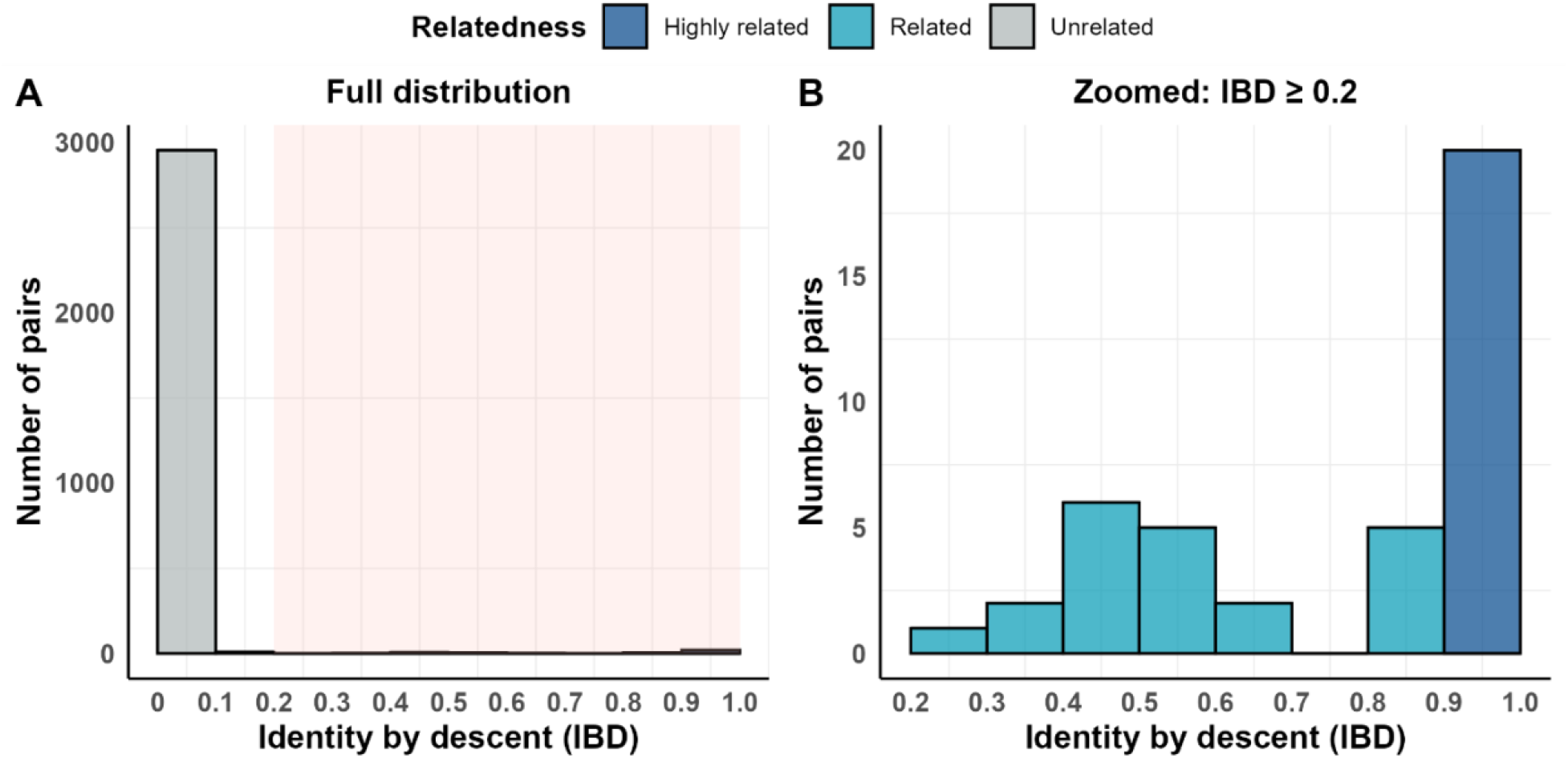
Distribution of Identity-By-Descent (IBD) values between *P. falciparum* genomes from community cases in the eastern Gambia. (A) Genetic relatedness between parasites was assessed by measuring IBD pairwise between 79 genomes, resulting in 3003 comparisons. (B) Zoom on values ≥ 0.2. Highly related and identical genomes (IBD > 0.9) are highlighted in dark blue.

### *De novo* SNP mutations inferred from highly related genomes

We compared identical or near-identical genomes (IBD > 0.9) and identified inferred *de novo* mutations as SNPs or indels present in one genome but absent from another otherwise identical genome. These mutations will have arisen since the identical genomes shared a common ancestor. A total of 20 highly related (IBD > 0.9) monoclonal genomes clustered into 8 IBD-defined clusters, each containing between two to four infection genome sequences (Table S3). Analysis was limited to the core genome, and known hypervariable genes were filtered out as these do not enable identification of new mutants using the methods here. Three clusters (12, 30 & 33; nine genomes in total), showed no *de novo* SNP (Figure 3A). In contrast, the remaining five clusters (total of 11 genomes) collectively harboured 43 *de novo* SNPs. The number of SNPs per cluster ranged from 1 to 18 SNPs, and these were distributed across all 14 chromosomes (Supplementary Figure 3). As expected, none of these SNPs were detected in any of the other monoclonal genomes (Table S3), supporting their recent origin within each cluster.

**Fig 3.**
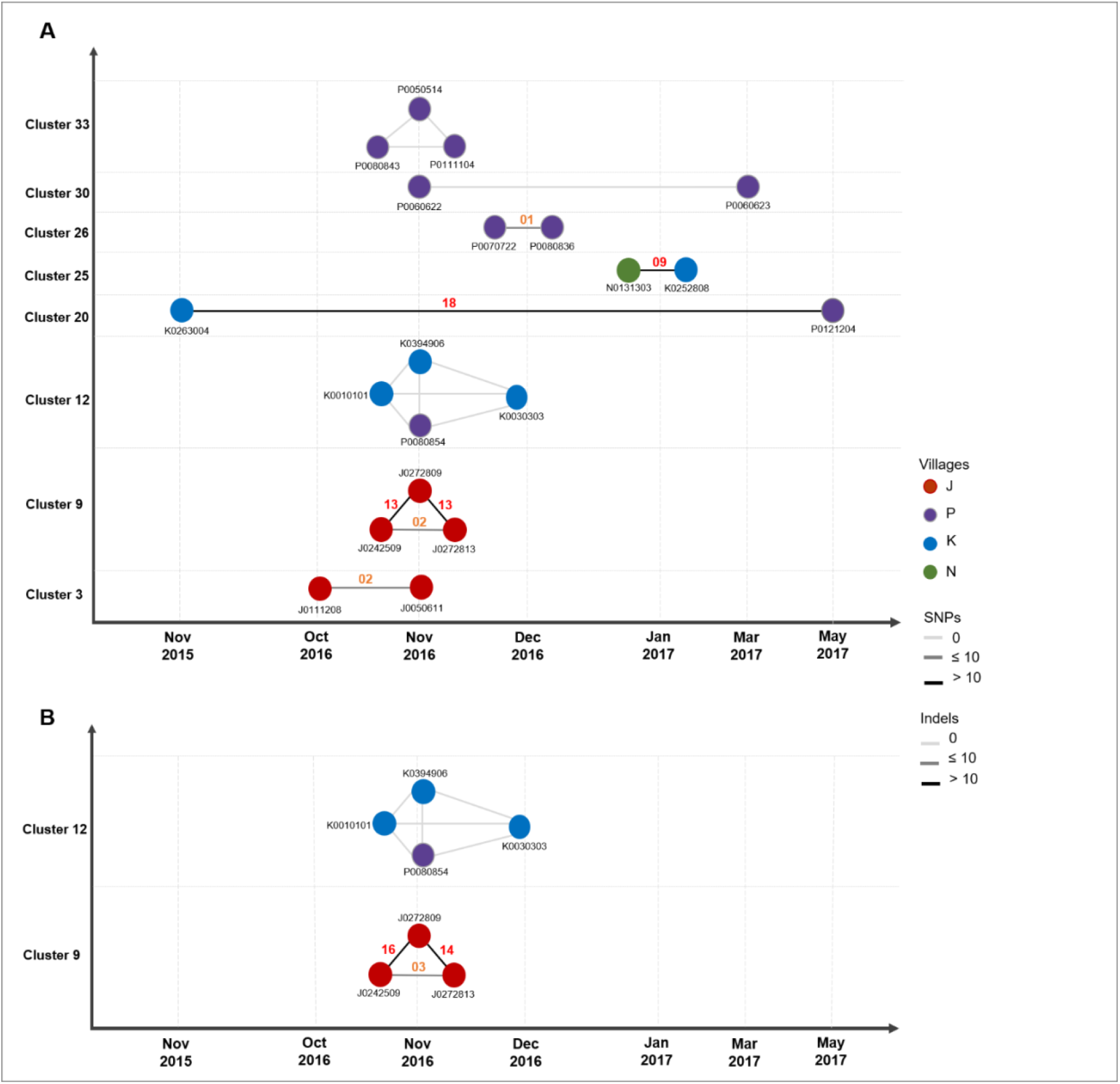
*De novo* mutations differentiating near-identical genomes. Clusters of near-identical genomes were defined as genomes with IBD >0.9. Each circle is a parasite genome, with the participant ID indicated; the first letter of the ID corresponds to the village of residence. The numbers on the plots indicate the count of *de novo* mutations differentiating pairs of genomes in IBD. (A) *De novo* SNP variants between near-identical genomes. Absence of *de novo* variants is indicated by light gray lines. (B) *De novo* short indel variants between near-identical genomes. Indel analysis was performed only in the IBD-defined genome groups (clusters 9 and 12) for which a long-read reference genome was available. The full list of *de novo* SNPs and indels is provided in Supplementary Tables 3 and 4.

Notably, cluster 25, comprising two genomes collected in the same month but from different villages, contained 9 *de novo* SNPs, suggesting that these parasites had diverged without outcrossing (Figure 3A).

Cluster 9 consisted of three genomes sampled within a single month from three individuals residing in nearby households. One genome (J0272809) carried 12 unique *de novo* SNPs absent in the other two genomes. By contrast, the latter pair (J0242509 & J0272813) differed only by two *de novo* SNPs. Given the narrow sampling interval, this pattern is consistent with J0272809 representing either a long-lasting chronic infection that accumulated mutations over time or the outcome of one or more rounds of transmission with self-fertilisation. In either case, these data indicate that J0242509 and J0272813 are more closely related to each other than to their most recent common ancestor with J0272809.

Another example of how genomic data can inform transmission patterns is provided by cluster 30. This cluster comprises two children from the same household. One child (J0020222) presented with symptomatic malaria in November 2016, whereas the second child (J0023023) harboured a prolonged asymptomatic infection from early December 2016 to May 2017. This latter infection was initially highly polyclonal, with the cluster 30 genotype only detected at later time points (March and May 2017) (Collins et al., 2022). Either J0020222 had transmitted to J002023 before receiving anti-malarial, either they acquired infections from the same mosquito, and the infection remained asymptomatic in J002023. This illustrates how combining genomic relatedness with longitudinal sampling can help disentangle recent transmission histories, even in the absence of detectable *de novo* mutations.

### *De novo* short indels identified using lineage-specific long-read reference genomes

We took advantage of two long-read sequencing genomes available for clusters 9 and 12 (Nyarko et al., 2025) and mapped Illumina reads against their respective long-read genome, followed by indel calling with GATK (Table S4). No *de novo* indels were detected in any of the 4 genomes from cluster 12, a result confirming the SNP analysis. All four genomes are therefore 100% identical, within the limit of our conservative analysis. In contrast, within the 3 genomes of cluster 9, 19 *de novo* short indels were identified (Figure 3B). Showing a parallel to the *de novo* SNP results, the genome sequence of infection J02728809 harbours 84.2% (16/19) of the *de novo* indels, setting it apart from the other two genomes within the cluster. Therefore the *de novo* indel analysis confirms the *de novo* SNP analysis and its interpretations.

### De novo *in vivo* mutations mirror *in vitro* mutation patterns

Our discovery of inferred *de novo* mutations is also a unique opportunity to study the mutation spectrum in *P. falciparum* genomes during natural infection conditions, as opposed to laboratory conditions. We further characterised the distribution and spectrum of the 43 *de novo* SNPs from community infections, and compared them to patterns observed in *in vitro* mutation accumulation experiments (Hamilton et al., 2016; McDew-White et al., 2019). Here, 29 (67%) *de novo* SNPs were located within exons; of these, 23 (79%) were Non-Synonymous and 6 (21%) Synonymous (Table 1). Although the proportion of synonymous mutations was lower than had been observed in a study of parasites in *in vitro* culture (31%), the difference was not significant (binomial test, p = 0.238). Non-synonymous mutations occurred across genes of diverse and mostly unknown functions. Notably, one *de novo* SNP introduced a premature stop codon in gene PF3D7_0301900 (unknown function), which would produce a truncated protein. That gene is essential for asexual development *in vitro* (Zhang et al., 2018). Such nonsense mutant alleles are very rare in wild parasite populations(Claessens et al., 2017), and it is unknown whether this apparent “natural knock-out” has a reduced fitness.

**Table 1.**
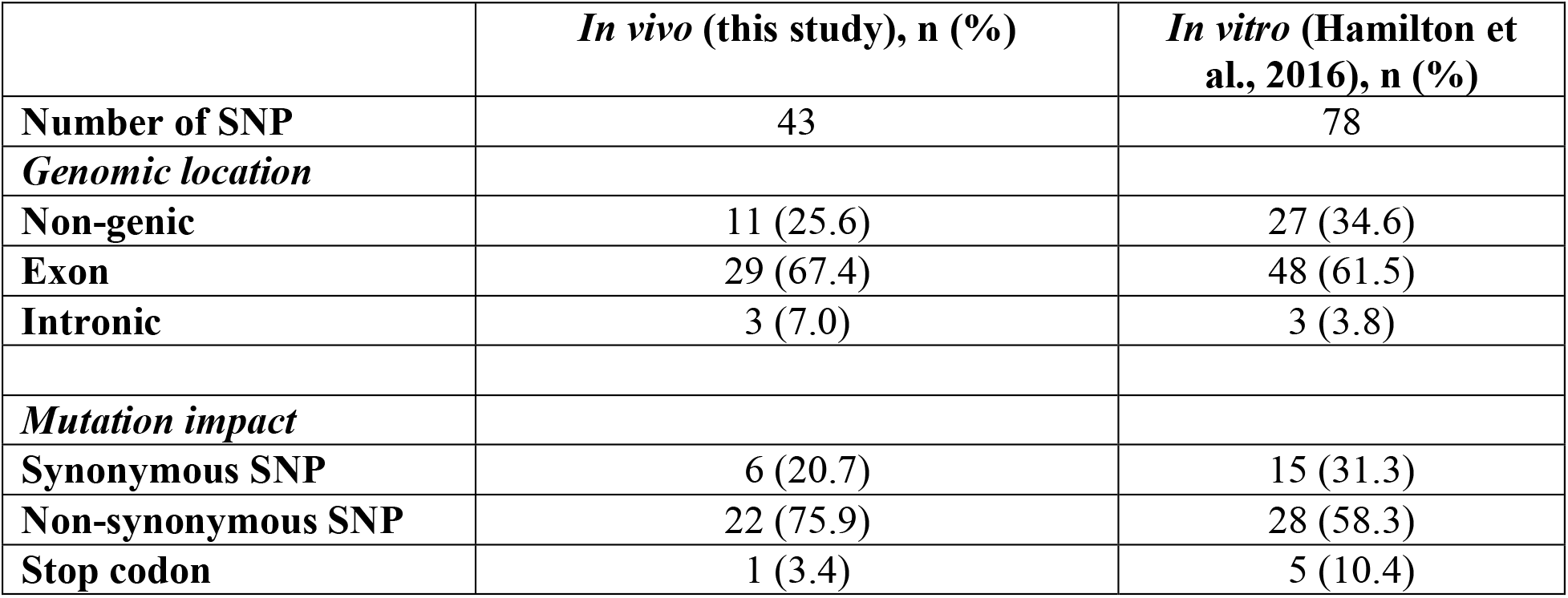
Genomic location and functional classification of *de novo* SNP mutations.

We next examined whether the strong mutational bias toward G:C → A:T transitions previously described *in vitro* could be detected *in vivo*, given that this bias is thought to underlie the extreme AT richness of the *P. falciparum* genome (80.6% in 3D7). If all nucleotide substitutions were equally likely, one would expect two transitions (A:T → G:C and G:C → A:T) for every four transversions, yielding a baseline transition-to-transversion (Ts:Tv) ratio of 0.5. Instead, we observed a Ts:Tv ratio of 3.3 (Table 2), substantially higher than the null expectation and exceeding the *in vitro* estimate of 1.13 (Hamilton et al., 2016). This enrichment in transitions was statistically significant (two-sided binomial test, p-value < 1.27e-10), consistent with a strong mutational bias operating *in vivo*.

**Table 2.** *de novo* SNP counts by substitution type in *P. falciparum* isolates.

| BPS | A:T→G:C | G:C→A:T | Ts | A:T→C:G | A:T→T:A | G:C→T:A | G:C→C:G | Tv | Ts:Tv |
| --- | --- | --- | --- | --- | --- | --- | --- | --- | --- |
| SNP counts | 17 | 16 | 33 | 1 | 4 | 4 | 1 | 10 | 3.3 |
| BPS/bp (1.10<sup>-6</sup>) | 0.91 | 3.54 |  | 0.05 | 0.21 | 0.88 | 0.22 |  |  |

To calculate Base Pair Substitution (BPS) rates for each substitution type, the raw numbers of BPS in each type (A:T→G:C, G:C→A:T, A:T→C:G, A:T→T:A, G:C→T:A, and G:C→C:G) were divided by the total number of A/T or G/C nucleotides in the *P. falciparum* core genome, taken as 18 779 800 bp and 4 520 200 bp, respectively (based on GC content of 19.36%), giving BPS/bp for each substitution type. Ts : Transition. Tv :Transversion.

Given the ∼80% AT composition of the genome, and assuming no mutational bias, A/T → G/C substitutions (i.e. A or T mutation to G or C) would be expected to occur approximately four times more frequently than G/C → A/T substitutions. However, we observed nearly equal numbers of the two classes, with 18 A/T→G/C substitution for 20 G/C→A/T substitution. Thus, G/C → A/T substitutions occurred 4.6-fold more frequently than expected under a neutral model. Under a simple mutation-equilibrium model (Sueoka, 1988), this skewed substitution rate predicts a mutational equilibrium genome composition of 82.15% AT, closely matching the observed AT content of 80.6%. These results indicate that the strong mutational bias toward G/C → A/T transitions is likely a major contributor to the extreme AT richness of the *P. falciparum* genome, although additional forces such as selection on codon usage may also contribute (Hershberg & Petrov, 2008, 2010).

*De novo* Indels ranged in size from 1 to 90 base pairs within cluster 9 (Table S4). Insertions accounted for 57.9% of all indels (Table 3), comparable proportions to those reported in *in vitro* mutation accumulation experiments (65.8%; chi-square test, χ^2^ = 0.4742, p = 0.4911) (Hamilton et al., 2016). The majority of indels (78.9%) were shorter than 3 bp in length, with median size of 1 bp for insertions and 2 bp for deletions (Table S4).

**Table 3.** Genomic location and classification of *de novo* short indels.

|  | *In vivo* (this study), n (%) | *In vitro* (Hamilton et al., 2016), n (%) |
| --- | --- | --- |
| **Number of Indels** | 19 | 164 |
| **Insertion** | 11 (57.9) | 108 (65.8) |
| **Deletion** | 8 (42.1) | 56 (34.1) |
| **Non-genic** | 14 (73.7) | 106 (64.6) |
| **Insertion** | 7 (50.0) | 73 (68.9) |
| **Deletion** | 7 (50.0) | 33 (31.1) |
| **Exon** | 3 (15.3) | 15 (9.1) |
| **Insertion** | 3 (100) | 3 (20.0) |
| **Deletion** | 0 (0.0) | 12 (80.0) |
| **Intronic** | 2 (10.5) | 43 (26.2) |
| **Insertion** | 1 (50.0) | 32 (74.4) |
| **Deletion** | 1 (50.0) | 11 (25.6) |

Most indels (73.7%) occurred in non-genic regions, significantly exceeding the proportion of non-genic sequence in the *P. falciparum* genome (41.9%; binomial test, *p* < 0.0081). Although more indels were observed in exons (n = 3) than in introns (n = 2), the total number of events is small and does not permit robust interpretation of this pattern.

Overall, the close concordance between *in vivo* and *in vitro* SNP and indel spectra indicates that the majority of *de novo* mutations observed in natural infections are neutral or nearly neutral. These results provide a reliable empirical baseline for interpreting fine-scale genetic relatedness and evolutionary dynamics among *P. falciparum* parasites in endemic populations.

## DISCUSSION

By combining identity-by-descent (IBD) analysis with the detection of de novo mutations, this study demonstrates how whole-genome sequencing can resolve genetic relationships among parasites circulating within a small community. Despite the small focused area of the study site, only a small proportion of pairwise comparisons between infections showed appreciable relatedness (IBD > 0.2). This suggests that, even in an area of declining transmission, sufficient parasite mixing, outcrossing and recombination persists to maintain a high level of genetic diversity. Similar patterns have been reported from other African regions transitioning toward pre-elimination, as population diversity remains high until transmission reaches very low levels (Atuh et al., 2022; Mobegi et al., 2014; Pringle et al., 2019; Sy et al., 2022). The rarity of highly related isolates illustrates why genotyping of few loci usually fails to resolve fine-scale relatedness in such settings, and underscores the value of WGS-based approaches (Mwesigwa et al., 2024).

If malaria control progresses in The Gambia, parasite prevalence is likely to decline further. As transmission decreases, parasite populations are predicted to become increasingly fragmented and dominated by a small number of persisting lineages. In such settings, the ability to accurately identify and track these residual lineages becomes critical to distinguish between local persistence and reintroduction, and to understand the origin of new infections or resurgences. The approach presented here, combining identity-by-descent analysis with the detection of *de novo* mutations, may clarify the exact nature of relatedness between such closely related parasites (Rowley et al., 2026; Tan et al., 2026). For example, genomic studies in South America have shown that malaria resurgences can be linked to transmission hotspots associated with human activities such as gold mining, which facilitate parasite spread across regions (Pacheco et al., 2020). Applying similar approaches could therefore enable the tracking of residual parasite lineages and help guide targeted interventions as countries move toward elimination.

Regions surrounding *crt* and *aat1* were disproportionately shared among unrelated genomes, consistent with previously documented selective sweeps in The Gambia (Amambua-Ngwa et al., 2023; Mobegi et al., 2014). The concordance with previous iHS-based analyses (Nwakanma et al., 2014) validates the capacity of IBD mapping to detect population-level WGS datasets, even when sample sizes are modest. Importantly, these selected regions were short and did not drive broad population structure, aligning with expectations in populations where drug use pressure is ongoing but recombination remains frequent (Guo et al., 2024).

*De novo* mutations are widely used to track transmission in viral and bacterial pathogens (Didelot et al., 2014; Eyre et al., 2013), but their application in malaria has been limited (Redmond et al., 2018). In this dataset, 3 out 8 clusters did not show any *de novo* mutation. This may reflect conservative filtering, but it also aligns with expectations: mutations arising within a host are often present at low frequency and may not reach fixation before transmission. Such variants are only captured in consensus genomes after transmission bottlenecks. Experimental infections suggest that these bottlenecks can be extremely tight, in some cases involving only a single or very few sporozoites (Pickford et al., 2021). This highlights the limitation of ‘bulk’ sequencing. In contrast, single-cell genomic methods have been shown to resolve within-host diversity and detect *de novo* mutations present at low frequency (Dia et al., 2021; Trevino et al., 2017).

The mutation spectrum observed in these natural infections resembles patterns previously described in laboratory mutation-accumulation experiments (Hamilton et al., 2016; McDew-White et al., 2019), and is consistent with broader evolutionary models in which mutational biases contribute substantially to genome composition (Hershberg & Petrov, 2010). In particular, we observed a strong transition bias and a pronounced excess of G:C→A:T substitutions, consistent with the mutational processes believed to contribute to the extreme AT richness of the *P*.

*falciparum* genome. Future studies could combine single-cell genomics with longitudinal sampling of chronic asymptomatic infections, allowing the accumulation and potential selection of adaptive mutations to be tracked over time.

In conclusion, this study demonstrates that integrating IBD analysis with *de novo* mutation detection provides a powerful framework for dissecting the fine-scale population structure of *P. falciparum*. As whole-genome sequencing becomes increasingly accessible, these approaches may help reconstruct parasite transmission networks, identify persistent reservoirs, and support malaria elimination strategies in regions approaching low transmission.

## Acknowledgements

The authors would like to thank the staff of MalariaGEN, Wellcome Sanger Institute Sample Management, Genotyping, Sequencing, and Informatics teams for their contribution. We thank Bahdja Boudoua. This work was funded by grants from the French National Research Agency (ANR 18-CE15-0009-01), CNRS Transversales, the Fondation pour la Recherche Médicale (EQU202303016290, EQU202203014741).

## Data Availability Statement

All sequencing data are available through the European Nucleotide Archive (ENA) under the accession numbers provided in the Supplementary Tables.

## Notes

### Competing Interest Statement

The authors have declared no competing interest.

