## Supplemental Figures for "Tracking malaria parasite lineages through *de novo* mutations in highly related *Plasmodium falciparum* genomes"

### Supplementary figures

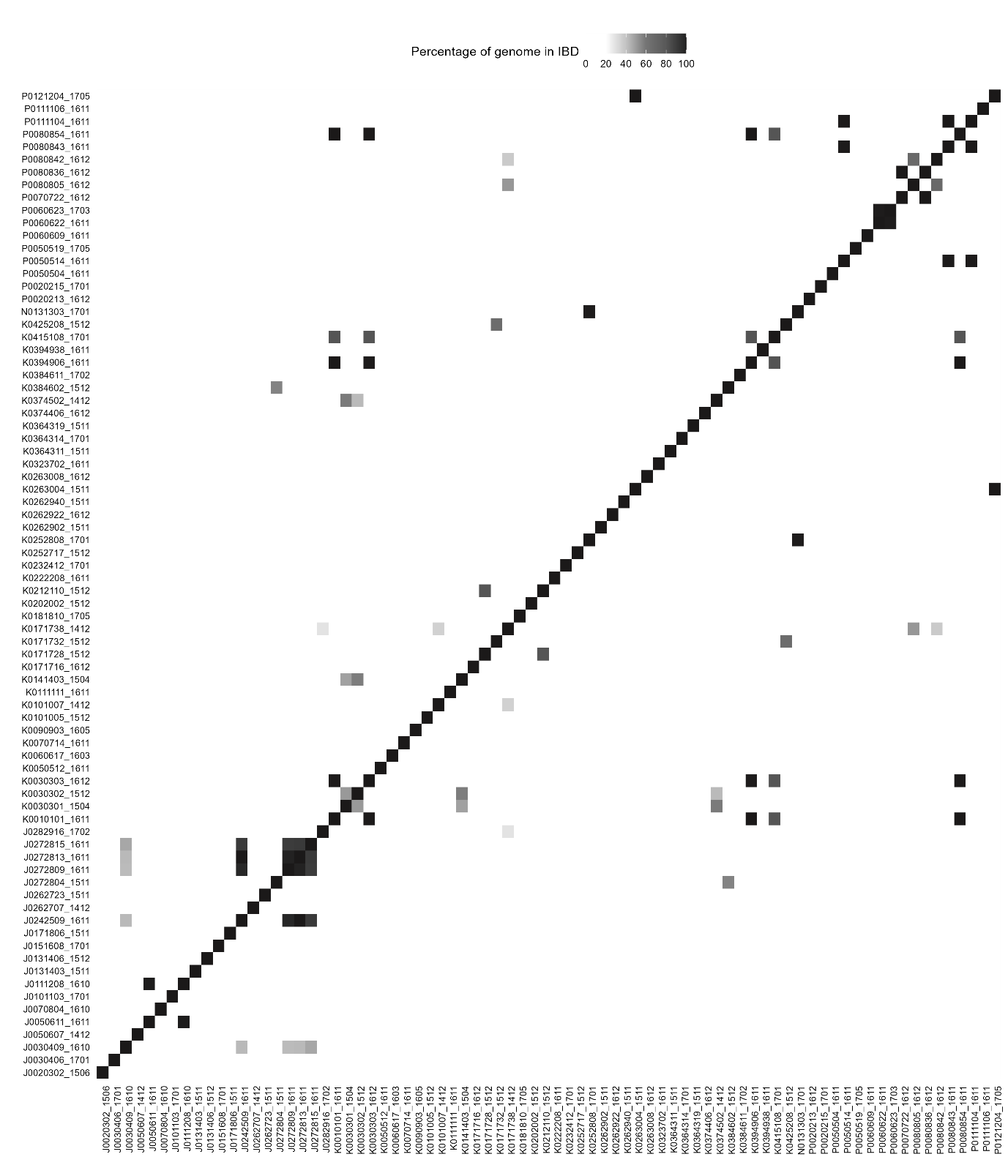

**Supp Figure 1: Percentage of genomic fragments in IBD between monoclonal genomes after merging identical (IBD > 0.9) genomes from the same individual.** All genomes were clustered according to their IBD values converted to a distance matrix.

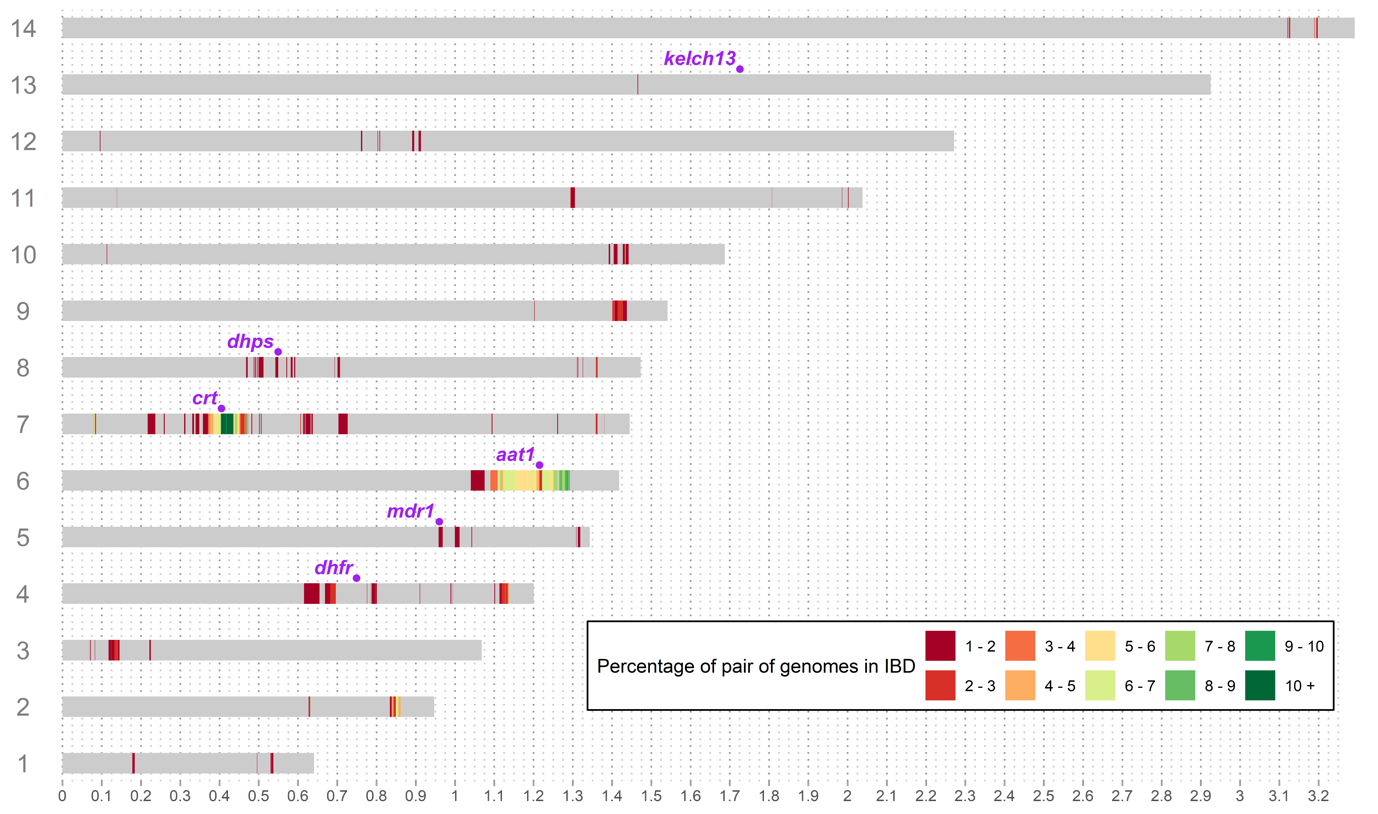

**Supp Figure 2. IBD proportion among unrelated *P. falciparum* genomes**.

We aimed to identify chromosomal regions under positive selection by examining IBD sharing among non-identical genomes (IBD < 0.5; 2,971 genome pairs). The percentage of genome pairs sharing a given DNA fragment in IBD is shown on a colour scale ranging from red (1–2% of pairs share the fragment) to green.

The 14 *P. falciparum* chromosomes are annotated with six genes known to be associated with reduced susceptibility to antimalarials: *aat1*, *crt*, *dhfr*, *dhps*, *kelch13*, and *mdr1*. Two regions with the highest levels of IBD sharing are centred on the *aat1* locus on chromosome 6 and the *crt* locus on chromosome 7. Strong linkage disequilibrium between these loci has been reported previously (Amambua-Ngwa et al., 2023). However, these regions are each smaller than 100 kb and therefore have minimal impact on genome-wide IBD estimates, as previously demonstrated in Southeast Asia (Guo et al., 2024).

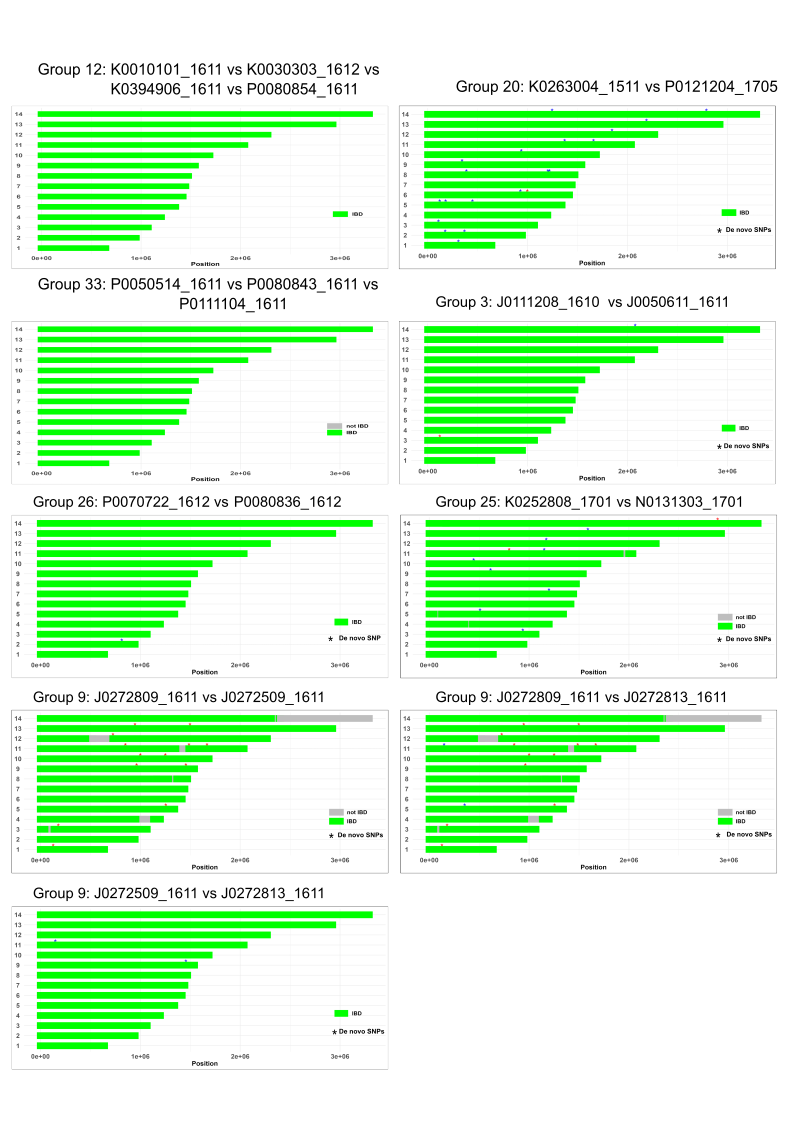

**Supp Fig 3. Distribution of *de novo* mutations distinguishing nearly identical genomes.** IBD clusters, comprising identical or nearly identical parasite genomes from 2 or more individuals, are clonal expansions. Green segments represent fragments of the chromosome that are identical by descent (IBD), indicating shared regions between two genomes. Only core genomes are represented here, i.e. hypervariable regions and subtelomeres were filtered out. Grey segments were not in IBD, they indicate outcrossing or unreliable calling regions, and were filtered out. Variants found within a clonal expansion represent *de novo* mutation in one or more genome. A red star indicates a *de novo* SNP unique to the first named isolate, while a blue star represents a *de novo* SNP exclusive to the second named isolate.
